# Visual Cortical Response Variability in Infants at High Familial Likelihood for Autism

**DOI:** 10.64898/2026.03.05.709374

**Authors:** Abigail Dickinson, Madison Booth, Scott Huberty, Declan Ryan, Alana Campbell, Jessica B. Girault, Neely C. Miller, Bonnie K. Lau, John M. Zempel, Sara Jane Webb, Jed T. Elison, Adrian KC Lee, Annette M. Estes, Stephen R. Dager, Heather C. Hazlett, Jason J. Wolff, Robert Schultz, Natasha Marrus, Alan C. Evans, Joseph Piven, John R. Pruett, Shafali S. Jeste, the IBIS Network

## Abstract

Visual processing undergoes rapid development in the first year of life, supporting the emergence of higher-order cognitive, language, and motor functions. Visual evoked potentials (VEPs) provide a non-invasive measure of visual system maturation that may shed light on heterogeneous developmental trajectories among infants at high familial likelihood for autism.

Infants with an older sibling with autism spectrum disorder (N = 177 at 6 months; N = 132 at 12 months) participated in the Infant Brain Imaging Study–Early Prediction (IBIS-EP) study. Pattern-reversal VEPs were recorded at 6 and 12 months, and developmental skills were assessed at 24 months using the Bayley Scales of Infant and Toddler Development (Bayley-4). VEP components (P1 and N1) were characterized by their amplitude and latency, as well as trial-to-trial variability in these measures. Associations with 24-month cognitive, language, and motor scores were examined using general linear models controlling for age, site, sex, and trial count.

Robust VEPs were observed at both time points, with age-appropriate morphology and expected developmental changes, including decreases in P1 latency and amplitude from 6 to 12 months. Greater trial-to-trial variability in P1 latency at both time points was associated with higher cognitive and language scores at 24 months. In contrast, conventional measures of mean P1 latency and amplitude were not associated with developmental outcomes.

These findings suggest that temporal variability in early visual responses may index adaptive sensory-circuit flexibility during a period of rapid experience-dependent development. VEP response-timing variability may therefore provide an early mechanistic marker of sensory-circuit organization relevant to later developmental trajectories.

**Research Highlights:**

- Greater trial-to-trial variability in visual cortical response timing was associated with higher cognitive and language scores in infants at high familial likelihood for autism.
- Conventional average VEP measures were unrelated to outcomes, suggesting temporal variability may more sensitively capture relevant neural-circuit differences in this population.
- Greater variability in early visual responses may reflect adaptive sensory-circuit flexibility during a critical period of neurodevelopment.
- VEP response-timing measures provide an early mechanistic window into sensory-circuit organization and its relation to later developmental trajectories.

## 1. Introduction

Identifying markers of atypical neurodevelopment before clinical diagnoses such as autism can be made is essential for understanding the mechanisms that shape developmental variability and informing earlier detection and intervention. The development of basic sensory systems may be particularly valuable in this context, as sensory systems are among the first functional neural networks to mature, and their maturation provides the foundation for higher-order processes, including attention, language, and social cognition (Braddick & Atkinson, 2011; Chorna et al., 2024; Siu & Murphy, 2018). The visual system undergoes rapid structural and functional refinement during the first year of life, supporting improvements in acuity, depth perception, motion tracking, and attentional control (M. H. Johnson, 1990). This period of visual circuit maturation provides the scaffolding for complex behaviors such as joint attention, face and object recognition, and the integration of visual input with motor planning and early language learning (M. H. Johnson et al., 2015). Disruptions in these early visual processes may limit opportunities to coordinate perception and action or to map visual information onto emerging communication systems, leading to cascading effects on broader developmental outcomes.

Visual evoked potentials (VEPs) provide a non-invasive, temporally precise, and scalable method for probing early sensory-cortical mechanisms. Recorded using electroencephalography (EEG), VEPs capture the brain’s time-locked response to visual stimuli and index neural signaling within occipital cortex and associated visual pathways. The VEP is a stereotyped EEG waveform characterized by distinct components, including an early negative deflection (N1) followed by a positive peak (P1), which together reflect synaptic transmission and postsynaptic activity in primary and extrastriate visual areas (Creel, 2019). Variation in the latency and amplitude of these early components is thought to index the maturation of visual circuits, including increased myelination, synaptic efficiency, and the excitatory–inhibitory balance (Siu & Murphy, 2018). Trial-to-trial variability in response timing also offers complementary information about the stability and reliability of neural signaling, reflecting the consistency of cortical recruitment and the precision of sensory encoding (Milne, 2011). Together, these features make VEPs a powerful tool for quantifying early circuit function and its role in supporting infant development.

Accumulating evidence links individual differences in these visual cortical responses to variation in developmental outcomes. For instance, P1 and N1 latencies have been associated with motor skills in young children (Otten et al., 2025) and with cognitive and psychomotor scores in infants with suspected developmental delays (Kim et al., 2018). Prospective longitudinal studies further indicate that early visual response properties are related to later developmental trajectories. For example, P1 latency at 3 months predicted cognitive scores at 18 months in infants exposed to gestational diabetes (Torres-Espínola et al., 2018), and P1 amplitude at 6 months was associated with motor, cognitive, and language outcomes at 27 months in children exposed to early adversity (Jensen et al., 2019). Beyond mean response properties, the trial-to-trial consistency of neural responses offers an additional window into circuit organization. Although variability in neural signals is sometimes treated as noise, converging evidence suggests that during early sensitive periods, greater variability may reflect a system that remains open to experience-dependent tuning, a property that may be functionally beneficial rather than detrimental (Deguire et al., 2023; Faisal et al., 2008).

Although VEP metrics have shown promise for characterizing early neural processing and later developmental outcomes in infants with elevated likelihood of altered neurodevelopment due to diverse biological and environmental factors (e.g., (Jensen et al., 2019; Torres-Espínola et al., 2018)), it remains unclear whether similar associations are present in infants at high familial likelihood (HL) for autism. HL infants, defined by having an older sibling with autism, have elevated rates of autism recurrence (∼20%; (Ozonoff et al., 2011, 2024)) and higher rates (∼30%) of broader neurodevelopmental delays (Charman et al., 2017; Messinger et al., 2013). As such, this group provides a unique opportunity to study mechanisms of atypical brain development, as they can be identified from birth, exhibit wide variability in developmental trajectories, and are at increased likelihood for atypical cognitive, language, and motor outcomes. Previous EEG research in HL infants has shown that early oscillatory activity is linked to later development, including associations between resting alpha power at 3 months and language outcomes at 12 months (Levin et al., 2017). However, it remains unclear whether evoked responses, such as VEPs, capture variability in early sensory-circuit maturation that is relevant to later developmental outcomes in this population.

In this study, we assessed whether individual differences in early visual cortical processing were associated with later developmental trajectories in infants with a high familial likelihood for autism enrolled in the Infant Brain Imaging Study (IBIS). IBIS is a multisite longitudinal cohort with standardized EEG acquisition and behavioral assessment across five sites, providing a large sample for examining this question. Specifically, we tested whether VEP metrics measured at 6 and 12 months were associated with cognitive, language, and motor outcomes at 24 months. Primary analyses focused on P1 amplitude, latency, and trial-to-trial variability in these measures, with corresponding N1 metrics examined in secondary analyses. We hypothesized that shorter P1 latencies would be associated with more favorable developmental outcomes and examined latency variability as a complementary marker of neural-circuit organization. The present report focuses on continuous developmental outcomes within this IBIS cohort; categorical autism outcomes will be examined once cohort accrual is complete and sufficient cases are available for adequately powered analyses.

## 2. Methods

### 2.1. Participants

Participants were enrolled in the Infant Brain Imaging Study–Early Prediction (IBIS-EP), a prospective multisite cohort study of infant siblings of children with autism. Data were collected across five sites: Washington University in St. Louis, the University of Washington in Seattle, Children’s Hospital of Philadelphia, the University of Minnesota, and the University of North Carolina at Chapel Hill. All infants had an older full sibling with a diagnosis of autism confirmed through medical records and standardized diagnostic and screening measures, including the Social Communication Questionnaire (SCQ; (Rutter, Bailey, et al., 2003)), and the Autism Diagnostic Interview–Revised (ADI-R; (Rutter, Le Couteur, et al., 2003)).

Eligibility criteria included: (1) gestational age > 36 weeks; (2) absence of medical or neurological conditions affecting growth, development, or cognition (e.g., seizure disorders) or significant sensory impairments (e.g., vision or hearing loss); (3) no known genetic syndromes associated with ASD; (4) no immediate family history of psychosis, schizophrenia, or bipolar disorder; (5) no contraindications for MRI; and (6) English as the primary language in the home. These criteria were assessed during a structured family history interview and are consistent with prior IBIS protocols (Emerson et al., 2017; Hazlett et al., 2017). All procedures were approved by a centralized institutional review board at Washington University in St. Louis. Written informed consent was obtained from a parent or guardian for each participant, in accordance with the Declaration of Helsinki.

EEG data were collected at 6 and 12 months of age, and behavioral assessments were completed at 24 months. The present analyses focused on participants with EEG at 6 and/or 12 months and available behavioral data at 24 months. The present analyses reflect data available as of January 2025; 24-month outcome visits will continue through early 2027. VEP data were collected from 180 infants at 6 months and 135 infants at 12 months. Of these recordings, usable VEP data were obtained from 177 of 180 infants at 6 months and 132 of 135 infants at 12 months. Corresponding 24-month behavioral data were available for 98 infants in the 6-month EEG sample and 97 infants in the 12-month EEG sample. Demographic characteristics of the final analytic sample are presented in Table 1.

**Table 1.** Sample Characteristics.

|  | 6 Month | 12 Month |
| --- | --- | --- |
| EEG Recorded, *n* | 180 | 135 |
| EEG Usable, *n* | 177 | 132 |
| Available 24-Month Behavioral Data, *n* | 98 | 97 |
| Age at EEG (months; *M (SD)*) | 6.79 (0.80) | 12.72 (0.71) |
| Sex (female; *n (%)*) | 53 (54.1%) | 49 (50.5%) |
| Race, *n (%)* |  |  |
| Asian | 5 (5.1%) | 1 (1.0%) |
| Black | 6 (6.1%) | 7 (7.2%) |
| Hispanic | 5 (5.1%) | 5 (5.2%) |
| More than one race | 12 (12.2%) | 8 (8.2%) |
| White | 70 (71.4%) | 76 (78.4%) |
| Ethnicity, *n (%)* |  |  |
| Hispanic | 24 (24.5%) | 26 (26.8%) |
| Non-Hispanic | 74 (75.5%) | 71 (73.2%) |
| Household Income, *n (%)* |  |  |
| <50K | 26 (26.5%) | 27 (27.8%) |
| 50K-150K | 44 (44.9%) | 42 (43.3%) |
| >150K | 23 (23.5%) | 25 (25.8%) |
| Unknown | 5 (5.1%) | 3 (3.1%) |
| Maternal Education, *n (%)* |  |  |
| High school or less | 11 (11.2%) | 11 (11.3%) |
| Some College or degree | 55 (56.1%) | 53 (54.6%) |
| Graduate Training | 31 (31.6%) | 32 (33.0%) |
| Unknown | 1 (1.0%) | 1 (1.0%) |
| Bayley-4 composite scores (24 months) |  |  |
| Cognitive | 96.94 (14.55) [55.00–135.00] | 96.60 (14.32) [55.00–135.00] |
| Language | 90.77 (22.46) [45.00–134.00] | 91.13 (22.15) [45.00–134.00] |
| Motor | 95.45 (12.32) [61.00–124.00] | 94.74 (12.25) [61.00–124.00] |

*Note.* Participant counts are not mutually exclusive, as some infants contributed EEG data at both 6 and 12 months. Specifically, 68 participants had EEG data at both time points and corresponding 24-month behavioral data. These participants are included in both columns.

### 2.2. Behavioral Assessments

At 24 months, participants included in the present analytic sample completed the Bayley Scales of Infant and Toddler Development, Fourth Edition (Bayley-4) (Bayley & Aylward, 2019), a standardized measure of early developmental functioning. The assessment was administered by a licensed clinical psychologist or trained developmental specialist, following standardized administration procedures. Age-corrected composite scores were derived for the Cognitive, Language, and Motor domains. Language scores combine receptive and expressive communication, and Motor scores combine fine- and gross-motor performance. Scores were based on each child’s chronological age at the time of assessment, in line with Bayley-4 scoring guidelines.

### 2.3. EEG Collection

Visual evoked potentials (VEPs) were collected as part of a broader EEG battery that included resting-state and auditory paradigms (for full details, see (Dickinson et al., 2024)). Acquisition procedures were standardized across IBIS-EP sites. Infants were seated on their caregiver’s lap in a dimly lit room, positioned 57 cm from a laptop screen displaying a standard pattern-reversal VEP stimulus. The stimulus consisted of a black-and-white checkerboard (36 × 36 checks; 21 × 21 cm) with a central red fixation point. The checkerboard had 99% contrast, a mean luminance of 80 cd/m^2^, and subtended approximately 21° of visual angle. The checkerboard underwent 160 phase reversals, occurring every 500 ms, with each phase reversal constituting one trial. Stimulus presentation was controlled by E-Prime (Psychology Software Tools), and phase reversals were marked using a screen-mounted photocell connected to the EEG amplifier. EEG was recorded using Net Amps amplifiers and high-density HydroCel Geodesic Sensor Nets (EGI) at a sampling rate of 500–1000 Hz (see (Dickinson et al., 2024) for site-specific collection details). Attention was monitored and documented by an experimenter throughout the session.

### 2.4. EEG Processing

EEG data were processed using EEGLAB (Delorme & Makeig, 2004) and custom MATLAB scripts (The MathWorks, Inc., Natick, MA). Continuous data were band-pass filtered (0.3–30 Hz, FIR) and any periods of inattention noted by the examiner were excluded. Data were then visually inspected to remove artifactual sections of data and channels. After removing artifacts, all datasets were interpolated to a standard 33-channel 10–10 montage (including O1, O2, Oz) and re-referenced to the average of all channels. Data were epoched −100 to 300 ms around stimulus onset (defined by photocell triggers) and baseline-corrected (−100 to 0 ms).

Analyses were restricted to conventional VEP electrode sites (O1, O2, Oz) overlying the occipital cortex, where visual responses are typically strongest (e.g., (Kovarski et al., 2019)). Trials were first screened for artifacts, and any trials containing eye movements, blinks, or voltages exceeding ±150 µV were excluded. The remaining artifact-free trials were then evaluated for VEP morphology. Only trials exhibiting a characteristic VEP waveform were retained, defined by a positive P1 peak (70–150 ms) preceded by a negative N1 deflection and a monotonic rise between the N1 and P1 components. Trials that did not meet these criteria were excluded. Examples of accepted and rejected trial morphologies are shown in Supplementary Figure 1.

Following artifact and morphology-based exclusions, participants contributed an average of 48.57 (SD = 17.72; Range = 15-98) trials at 6 months and 42.24 (SD = 13.93; Range = 19-97) at 12 months. The number of usable trials was included as a covariate in all analyses, as described below.

For each occipital channel, quality-controlled trials were averaged to yield a mean VEP, from which the latency and amplitude of the P1 and N1 components were extracted. Component amplitudes were quantified relative to the pre-stimulus baseline. Trial-level amplitudes and latencies were used to compute intra-individual variability, indexed by the median absolute deviation (MAD) across trials for both amplitude and latency

(Kovarski et al., 2019; Milne, 2011). P1 and N1 metrics were extracted separately at O1, Oz, and O2 and then averaged across the three channels for each participant. Because the primary aim was to quantify trial-to-trial variability, analyses focused on the P1 component, which is the largest and most reliably identified early visual response at the single-trial level. Primary analyses therefore examined P1 latency, amplitude, and trial-to-trial variability in both measures, while the corresponding N1 metrics were evaluated in secondary analyses. Descriptive statistics (mean, standard deviation, and range) for all P1 and N1 metrics at each time point are presented in Table 2.

**Table 2.**
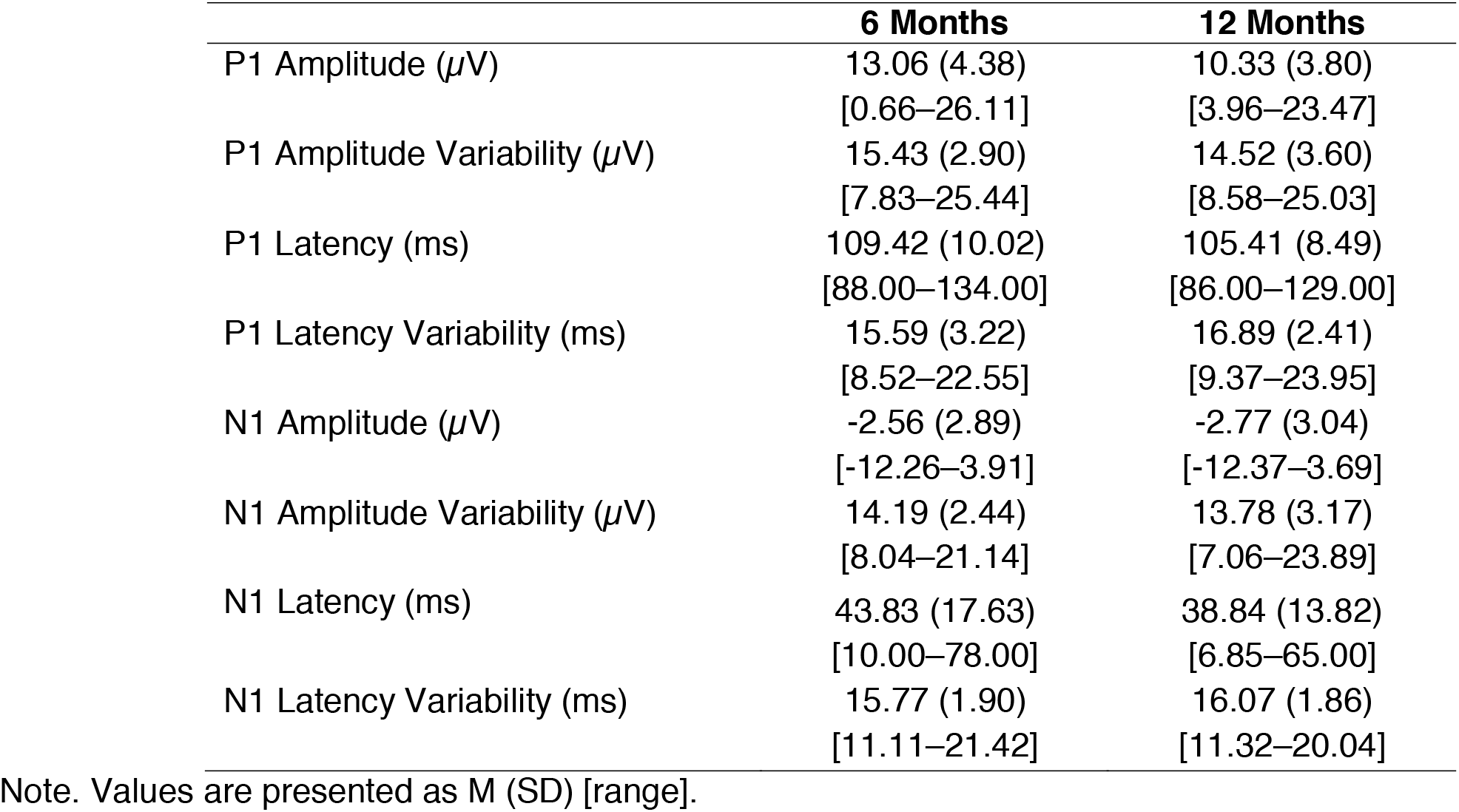
VEP Metric Values.

|  | **6 Months** | **12 Months** |
| --- | --- | --- |
| P1 Amplitude ($\mu V$) | 13.06 (4.38)<br>[0.66–26.11] | 10.33 (3.80)<br>[3.96–23.47] |
| P1 Amplitude Variability ($\mu V$) | 15.43 (2.90)<br>[7.83–25.44] | 14.52 (3.60)<br>[8.58–25.03] |
| P1 Latency (ms) | 109.42 (10.02)<br>[88.00–134.00] | 105.41 (8.49)<br>[86.00–129.00] |
| P1 Latency Variability (ms) | 15.59 (3.22)<br>[8.52–22.55] | 16.89 (2.41)<br>[9.37–23.95] |
| N1 Amplitude ($\mu V$) | -2.56 (2.89)<br>[-12.26–3.91] | -2.77 (3.04)<br>[-12.37–3.69] |
| N1 Amplitude Variability ($\mu V$) | 14.19 (2.44)<br>[8.04–21.14] | 13.78 (3.17)<br>[7.06–23.89] |
| N1 Latency (ms) | 43.83 (17.63)<br>[10.00–78.00] | 38.84 (13.82)<br>[6.85–65.00] |
| N1 Latency Variability (ms) | 15.77 (1.90)<br>[11.11–21.42] | 16.07 (1.86)<br>[11.32–20.04] |

Note. Values are presented as M (SD) [range].

**Figure 1.**
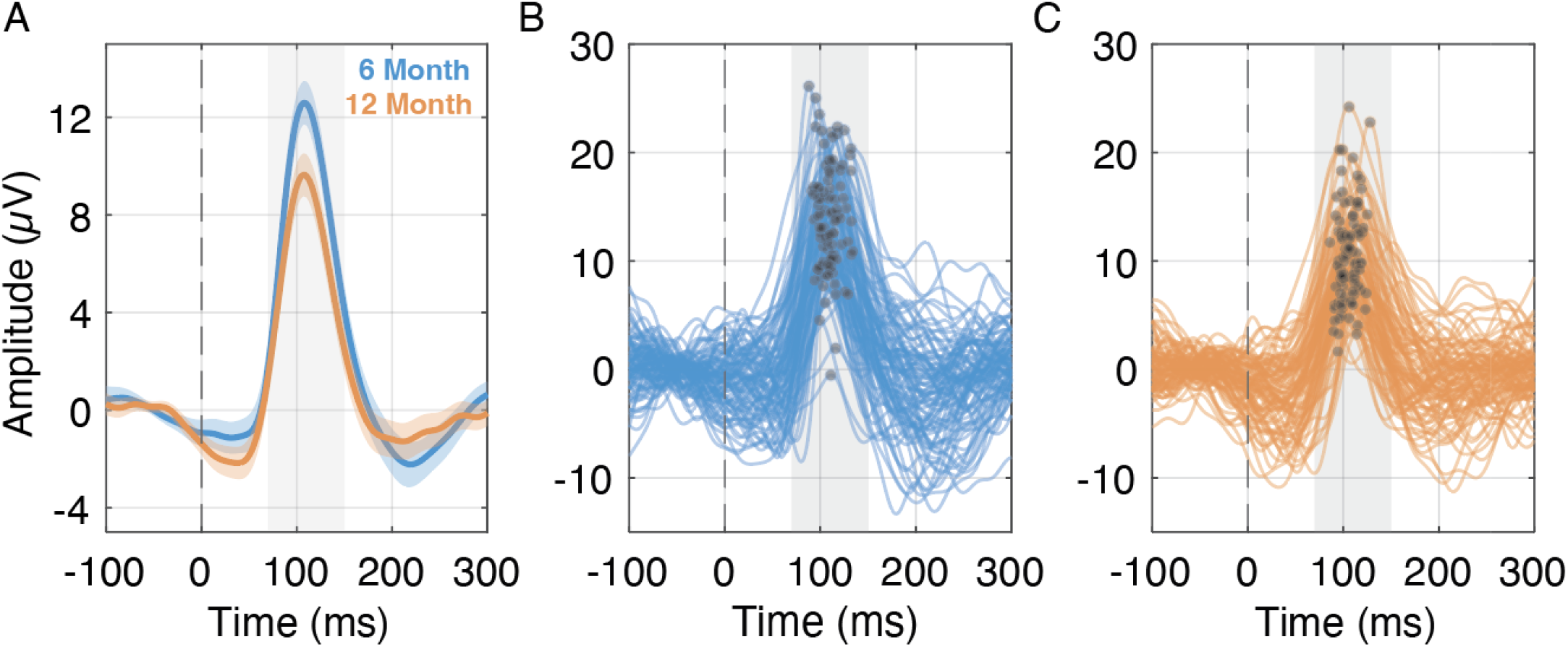
A) Grand average pattern-reversal visual evoked potentials (VEPs) at 6 months (blue) and 12 months (orange). Solid lines represent the mean VEP waveform across participants; shaded regions depict the 95% confidence interval. B–C) Individual VEP waveforms from participants at 6 months (B) and 12 months (C). Each line reflects the average waveform for a single participant. P1 components are marked with filled circles within the typical P1 latency window.

**Figure 2.**
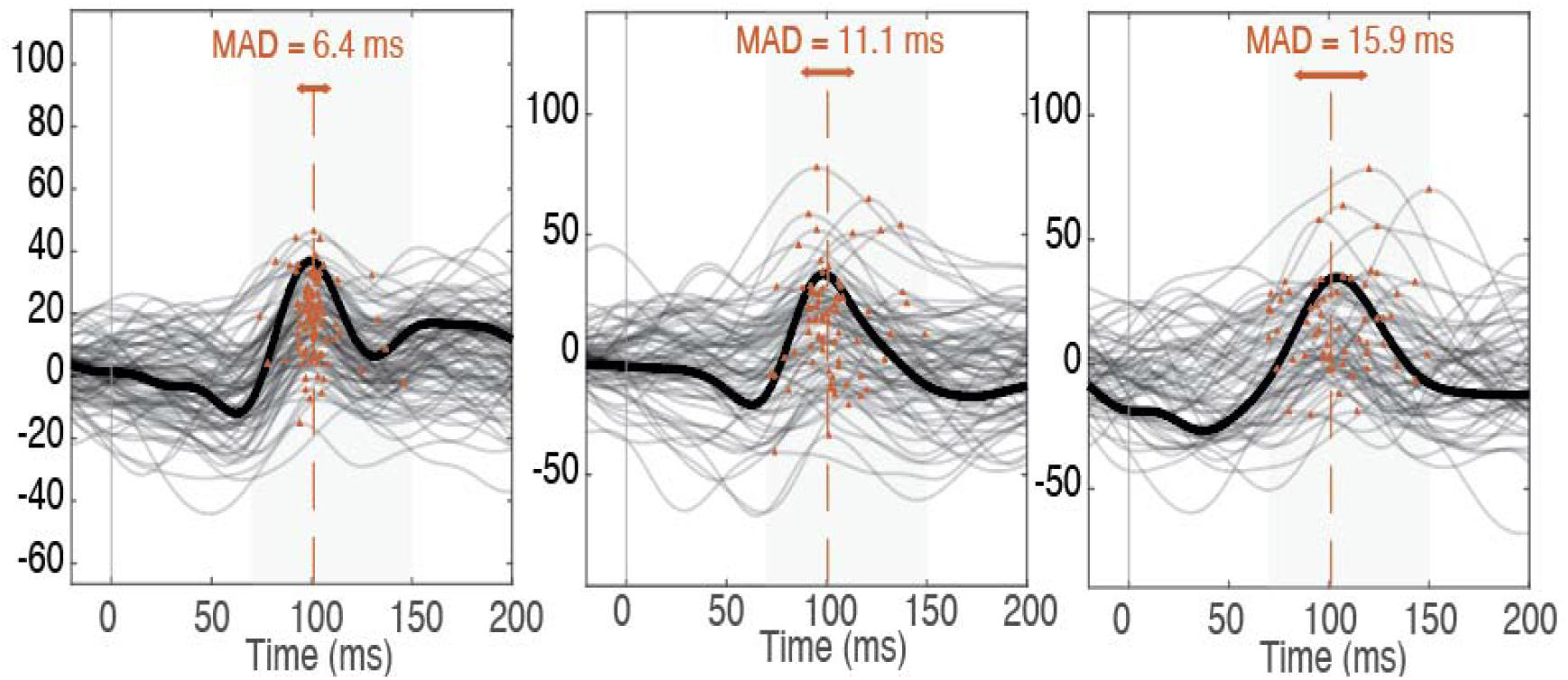
Examples of trial-level P1 identification in individual participants, illustrating differences in P1 latency variability. Each trace represents an individual trial, and the identified P1 component on each trial is marked with a red dot. Intra-individual P1 latency variability was quantified as the median absolute deviation (MAD) of P1 latencies across trials. Panels A–C show participants with low (A), intermediate (B), and high (C) trial-to-trial P1 latency variability. Although their average VEP waveforms may appear similar, the timing of the identified P1 components across individual trials differs substantially.

### 2.5. Statistical Analyses

Statistical analyses were conducted in MATLAB (The MathWorks, Inc., Natick, MA) and R. Four separate linear mixed-effects models were used to characterize developmental changes in each P1 metric (latency, amplitude, latency variability, and amplitude variability) from 6 to 12 months. Each model included time point as a fixed effect and participant as a random intercept to account for repeated measurements. Site, sex, and usable trial count were included as fixed-effect covariates. Developmental-change analyses included all participants with usable VEP data at one or both EEG assessments. VEP–behavior association analyses, described below, were restricted to participants with corresponding 24-month Bayley-4 data.

General linear models examined associations between VEP metrics measured at 6 and 12 months and cognitive, language, and motor outcomes assessed at 24 months. For each EEG time point and each developmental outcome, a single model included all four VEP metrics simultaneously (P1 amplitude, P1 latency, P1 latency variability, and P1 amplitude variability), allowing the unique contribution of each VEP metric to be estimated while adjusting for the others. This yielded three models per time point (one per outcome), for a total of six models. All models additionally included age at EEG acquisition (in months), site (five-level categorical variable), sex, and the number of usable trials as covariates. Continuous predictors were standardized prior to model fitting so that coefficients are expressed in standard-deviation units and are comparable across metrics. Variance inflation factors were examined to assess whether collinearity among the VEP metrics compromised coefficient estimates.

False discovery rate (FDR) correction using the Benjamini–Hochberg procedure was applied separately within the 6-month and 12-month model families to adjust for multiple comparisons, with each family comprising all VEP metric-by-outcome tests at that time point (four metrics × three outcomes). Table 3 reports standardized regression coefficients, two-sided p-values, and FDR-corrected q-values for the VEP predictors of interest, with associations meeting q ≤ .05 indicated in bold. Full regression outputs for all models, including covariate coefficients and variance inflation factors, are provided in Supplementary Tables S1–S2. Supplementary analyses examine associations between N1 components, using the same model specification, covariates, and FDR correction as for P1 components. These results are presented in Supplementary Table S3.

**Table 3.** Associations between EEG metrics and developmental outcomes at 24 months. Values represent regression coefficients (b) and corresponding *p* values for the EEG predictors of interest from each model. For each developmental outcome and EEG time point, all four P1 metrics were entered simultaneously into a single model. Significant associations that met the FDR-corrected significance threshold (*q* ≤ .05) are bolded and marked with an asterisk. Complete model results, including coefficients for all covariates, are provided in Supplementary Tables S1-S2.

| VEP Metric | 24-Month Behavioral Composite Scores |  |  |
| --- | --- | --- | --- |
|  | Cognitive | Language | Motor |
|  | *b* (*p*; *q*) | *b* (*p*; *q*) | *b* (*p*; *q*) |
| *6 Months* |  |  |  |
| P1 Amplitude (μV) | 1.47 (.456; .650) | 0.90 (.764; .833) | -1.76 (.291; .499) |
| P1 Amplitude Variability (μV) | -2.97 (.099; .198) | -5.74 (.038; .092) | -3.37 (.029; .092) |
| P1 Latency (ms) | 0.17 (.917; .917) | 1.06 (.676; .811) | -0.98 (.487; .650) |
| P1 Latency Variability (ms) | **7.34 (<.001; .005)\*** | **8.81 (.005; .028)\*** | 3.55 (.038; .092) |
| *12 Months* |  |  |  |
| P1 Amplitude (μV) | -1.75 (.410; .697) | 2.02 (.523; .697) | -1.03 (.581; .697) |
| P1 Amplitude Variability (μV) | 1.30 (.525; .697) | -0.82 (.789; .789) | 0.55 (.760; .789) |
| P1 Latency (ms) | 2.87 (.076; .229) | 2.21 (.358; .697) | 2.14 (.136; .325) |
| P1 Latency Variability (ms) | **4.54 (.008; .050)\*** | **6.95 (.007; .050)\*** | 2.79 (.065; .229) |

## 3. Results

### 3.1. Developmental Changes in P1 Metrics

Linear mixed-effects models accounting for repeated measurements, site, sex, and usable trial count indicated significant developmental changes in all four P1 metrics from 6 to 12 months. Mean P1 latency decreased from 109.42 ms (SD = 10.02) at 6 months to 105.41 ms (SD = 8.49) at 12 months, b = −4.22, SE = 1.19, t(187) = −3.54, p < .001. Mean P1 amplitude also decreased from 13.06 µV (SD = 4.38) at 6 months to 10.33 µV (SD = 3.80) at 12 months, b = −2.56, SE = 0.48, t(187) = −5.33, p < .001. In contrast, trial-to-trial variability in P1 latency increased from 15.59 ms (SD = 3.22) at 6 months to 16.89 ms (SD = 2.41) at 12 months, b = 0.87, SE = 0.32, t(187) = 2.70, p = .008. P1 amplitude variability decreased from 15.43 µV (SD = 2.90) at 6 months to 14.52 µV (SD = 3.60) at 12 months, b = −1.10, SE = 0.45, t(187) = −2.45, p = .015.

### 3.2. Associations between VEP metrics and 24-month developmental outcomes

General linear models examined associations between VEP metrics at 6 and 12 months and cognitive, language, and motor outcomes at 24 months. At both time points, greater trial-to-trial variability in P1 latency was associated with higher cognitive and language scores. Specifically, greater P1 latency variability at 6 months was associated with higher cognitive (b = 7.34, p < .001) and language (b = 8.81, p = .005) composite scores at 24 months, and both associations met the FDR-corrected significance threshold (q = .005 and q = .028, respectively). A nominal association with motor scores did not survive FDR correction (b = 3.55, p = .038, q = .092). At 12 months, greater P1 latency variability was similarly associated with higher cognitive (b = 4.54, p = .008, q = .050) and language (b = 6.95, p = .007, q = .050) scores. Both associations met the prespecified FDR threshold of q ≤ .05.

In contrast, mean P1 latency, mean P1 amplitude, and P1 amplitude variability were not significantly associated with developmental outcomes at either time point after FDR correction (all q ≥ .092). Detailed model estimates for each outcome and EEG metric are reported in Supplementary Tables S1 (6-month EEG) and S2 (12-month EEG).

Secondary analyses of N1 metrics showed a partially similar but weaker pattern (Supplementary Table S3). At 6 months, greater N1 latency variability was positively associated with cognitive (b = 3.54, p = .029) and motor scores (b = 3.66, p = .007) and showed a similar positive pattern for language scores (b = 4.84, p = .052).However, only the association with motor scores met the FDR-corrected significance threshold (q = .041); the cognitive and language associations did not (q = .115 and q = .125, respectively). N1 latency variability at 12 months was not associated with any developmental outcome after correction for multiple comparisons. Mean N1 latency and amplitude were also unrelated to developmental outcomes after FDR correction. N1 amplitude variability was not associated with developmental outcomes, except for a negative association with motor scores at 6 months (b = −3.96, p = .002, q = .025).

**Figure 3.**
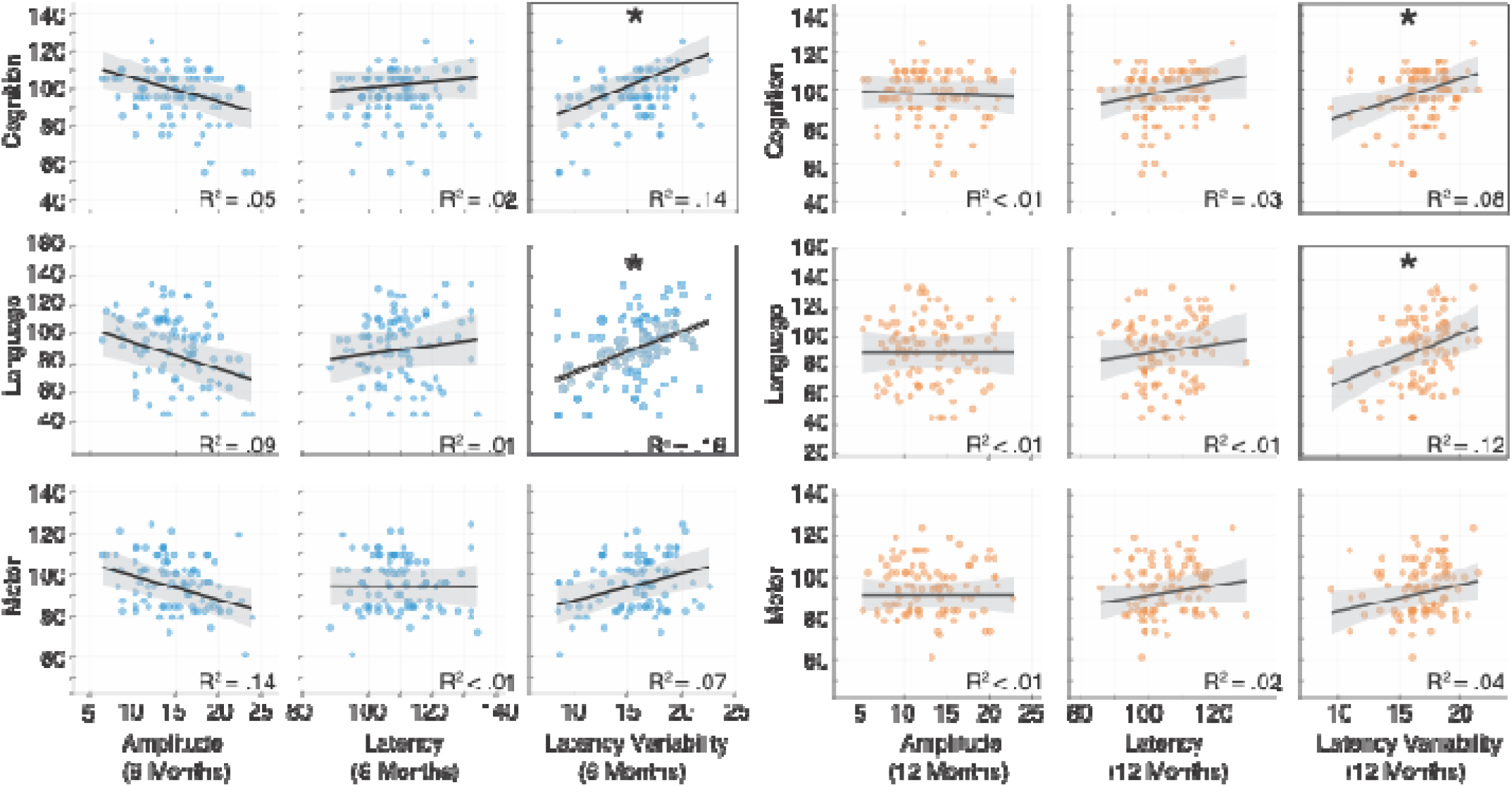
Scatterplots illustrating associations between EEG metrics (P1 latency, P1 latency variability, and P1 amplitude) measured at 6 months (blue) and 12 months (orange) and developmental outcomes at 24 months. The plotted regression lines and R^2^ values represent unadjusted bivariate associations and are provided for visualization; statistical significance was determined from the fully adjusted multivariable models. Asterisks identify associations that met the FDR-corrected significance threshold (*q* ≤ .05) in the fully adjusted multivariable models.

## 4. Discussion

This study demonstrates that trial-to-trial variability in the timing of early visual cortical responses is associated with later cognitive and language development in HL infants. Greater P1 latency variability at both 6 and 12 months was consistently associated with higher cognitive and language scores at 24 months, whereas conventional VEP measures of mean latency and amplitude were not. These findings suggest that temporal variability may capture developmentally relevant aspects of early visual-circuit organization that are not evident in conventional VEP averages.

Across the cohort, VEPs showed age-appropriate morphology and expected developmental change. P1 latency and amplitude both decreased from 6 to 12 months, consistent with ongoing myelination and increasing efficiency within visual pathways during the first year of life (Kovarski et al., 2019; MCculloch, 2013; Skoczenski & Norcia, 2002). Despite their sensitivity to age-related maturation, these mean measures were not associated with later cognitive, language, or motor outcomes. Prior studies have reported mixed associations between conventional infant VEP measures and later development, including links with P1 latency in some populations and P1 amplitude in others (Jensen et al., 2019; Otten et al., 2025; Torres-Espínola et al., 2018).

Neural response variability is a hallmark of early brain development. Newborns exhibit substantially greater VEP variability than older infants (Benavente et al., 2005), and neural variability generally declines with maturation (Hultsch et al., 2011). In this context, the association between greater P1 latency variability and higher later developmental scores suggests that variability may not simply reflect inefficient or unreliable signaling. Instead, it may index sensory circuits that remain flexible and responsive to experience during a period of rapid organization. A partially similar but weaker pattern was observed for N1 latency variability at 6 months, although only its association with motor outcomes remained significant after correction for multiple comparisons. Associations involving latency variability were also descriptively stronger earlier in infancy: P1 latency variability was related to cognitive and language outcomes at both ages, whereas N1 latency variability was related to outcomes only at 6 months. This pattern raises the possibility that neural variability is particularly informative during early sensitive periods, when sensory circuits remain highly plastic.

Greater response variability may therefore reflect an actively organizing visual system that has not yet fully stabilized. By remaining responsive to sensory experience during a critical period of learning, visual pathways may support the development of downstream cognitive, language, motor, and social functions (Braddick & Atkinson, 2011; Chorna et al., 2024; Siu & Murphy, 2018). This interpretation is consistent with cascading models of neurodevelopment, in which the early organization of sensory circuits provides a foundation for the emergence of higher-order abilities (S. P. Johnson, 2011; Karmiloff-Smith, 1998). Similar developmental accounts have been proposed for autism, emphasizing how early differences in basic sensory and neural systems may shape later-emerging phenotypes (Girault, 2025; Piven et al., 2017).

Converging evidence from other imaging modalities supports a functional role for neural variability. Greater moment-to-moment BOLD variability has been associated with neural flexibility and cognitive performance (Armbruster-Genç et al., 2016; Nomi et al., 2017), while recent EEG work suggests that infants with more differentiated neural responses to changing sensory input show better adaptive functioning at 4 years of age (Deguire et al., 2023). Together with the present findings, this body of work suggests that elevated neural variability early in development may reflect prolonged plasticity and exploratory circuit tuning that benefits learning. This interpretation is also consistent with prior structural MRI findings suggesting that early maturation of visual and associated white-matter pathways is important for later development. Across separate infant samples, including a prior IBIS cohort, individual differences in the development of these pathways have been linked to variation in later cognitive and language outcomes (Girault et al., 2022; Swanson et al., 2017). Taken together, these structural and functional findings suggest that early developmental advantage may arise not from rapid stabilization of neural systems, but from extended windows of plasticity that allow circuits to be shaped by experience. A critical next step will be to examine white-matter development in the same infants studied here to determine whether greater inter-trial variability in visual cortical responses is associated with more gradual or protracted white-matter maturation.

The present findings should also be considered alongside evidence of elevated neural-response variability in older autistic children and adults. In these populations, increased trial-to-trial variability in evoked responses has often been interpreted as reflecting unreliable or inefficient neural signaling (Dinstein et al., 2012; Dong et al., 2024; Milne, 2011). The contrasting interpretation proposed here highlights the importance of developmental context. Variability may reflect adaptive flexibility during early sensitive periods but unstable or dysregulated processing later in development. Longitudinal studies are needed to determine how variability changes across childhood, whether early variability is associated with later autism outcomes, and how infant response patterns relate to those observed in older autistic individuals.

Several limitations should be considered. Although this study included a large, well-characterized cohort, analyses were limited to infants at high familial likelihood for autism. While this focus is important for understanding early neural markers that may elucidate later developmental variability within this highly heterogeneous group, future work should examine HL infants alongside infants from the general population to determine whether the observed associations generalize across likelihood groups or reflect patterns specific to familial risk. In addition, the present analyses were designed to characterize continuous variation in developmental outcomes within the HL cohort rather than categorical diagnostic endpoints. Future analyses will test whether early differences in sensory-cortical processing are also associated with later categorical diagnostic outcomes once the full sample, consistent with pre-specified power targets, is available. Finally, it should be noted that our sample was disproportionately White, and parents were on average highly educated and relatively affluent (Table 1). This limits the generalizability of our findings, as socioeconomic and sociodemographic factors can influence both early neurodevelopment and developmental assessment.Replication in more diverse and representative samples is an important priority.

In summary, this study demonstrates that trial-to-trial variability in the timing of early visual cortical responses is associated with later cognitive and language development in HL infants. Rather than reflecting neural noise, temporal variability may capture subtle aspects of early sensory circuit maturation that are not apparent in conventional measures of average VEP amplitude or latency. By characterizing these within-individual dynamics, trial-to-trial variability may provide a mechanistic window into the evolving organization of visual cortical circuits and how early sensory processing scaffolds subsequent developmental trajectories.

