## Supplementary Figure 1 for "Visual Cortical Response Variability in Infants at High Familial Likelihood for Autism"

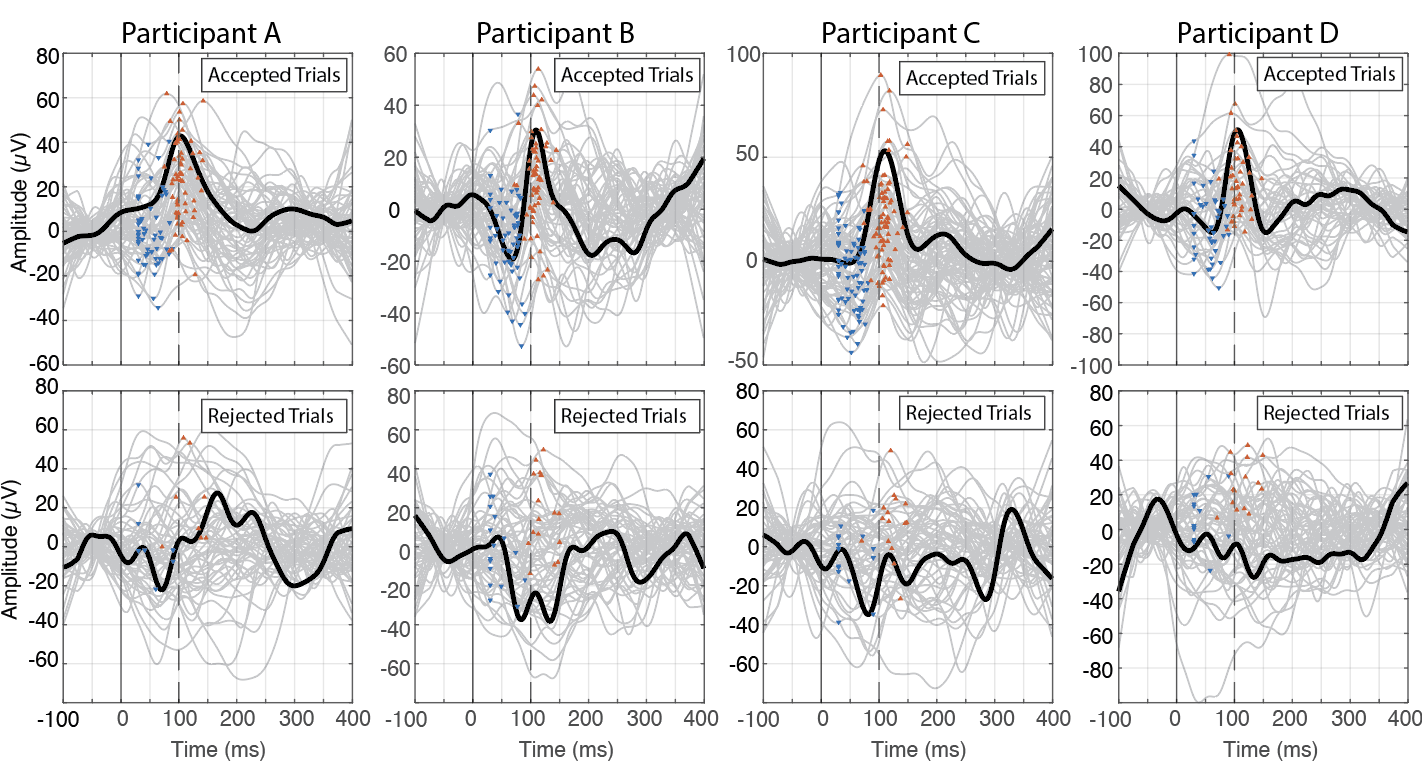
**Supplementary Figure 1.** Trial-level inclusion criteria and examples of retained versus excluded VEP trials. Trials were retained only if the peak amplitude did not exceed ±150 µV and the waveform exhibited a canonical N1–P1 morphology, which was defined as a positive deflection (P1; 70–150 ms) preceded by a negative deflection (N1), with a monotonic rise between the two. Panels A–D show four representative participants. Upper panels display retained trials (gray lines) with identified P1 (red triangles) and N1 (blue triangles); the black line indicates the average across trials. Lower panels show excluded trials that were rejected due to excessive amplitude, absent or mis-ordered peaks, or atypical morphology.
